# Heritability, selection, and the response to selection in the presence of phenotypic measurement error: effects, cures, and the role of repeated measurements

**DOI:** 10.1101/247189

**Authors:** Erica Ponzi, Lukas F. Keller, Timothée Bonnet, Stefanie Muff

**Author notes:** **Corresponding author:** Stefanie Muff, Department of Evolutionary Biology and Environmental Studies, University of Zürich, Winterthurerstrasse 190, 8057 Zürich, Switerland Department of Biostatistics, Epidemiology, Biostatistics and Prevention Institute, University of Zürich, Hirschengraben 84, 8001 Zürich, Switzerland.

## Abstract

Quantitative genetic analyses require extensive measurements of phenotypic traits, a task that is often not trivial, especially in wild populations. On top of instrumental measurement error, some traits may undergo transient (*i.e*. non-persistent) fluctuations that are biologically irrelevant for selection processes. These two sources of variability, which we denote here as measurement error in a broad sense, are possible causes for bias in the estimation of quantitative genetic parameters. We illustrate how in a continuous trait transient effects with a classical measurement error structure may bias estimates of heritability, selection gradients, and the predicted response to selection. We propose strategies to obtain unbiased estimates with the help of repeated measurements taken at an appropriate temporal scale. However, the fact that in quantitative genetic analyses repeated measurements are also used to isolate permanent environmental instead of transient effects, requires a re-assessment of the information content of repeated measurements. To do so, we propose to distinguish “short-term” from “long-term” repeats, where the former capture transient variability and the latter the permanent effects. We show how the inclusion of the corresponding variance components in quantitative genetic models yields unbiased estimates of all quantities of interest, and we illustrate the application of the method to data from a Swiss snow vole population.

## Introduction

Quantitative genetic methods have become increasingly popular for the study of natural populations in the last decades, and they now provide powerful tools to investigate the inheritance of characters, and to understand and predict evolutionary change of phenotypic traits (Falconer and Mackay, 1996; Lynch and Walsh, 1998; Charmantier et al., 2014). At its core, quantitative genetics is a statistical approach that decomposes the observed phenotype *P* into the sum of additive genetic effects *A* and a residual component *R*, so that *P* = *A* + *R*. For simplicity, non-additive genetic effects, such as dominance and epistatic effects, are ignored throughout this paper, thus the residual component can be thought of as the sum of all environmental effects. This basic model can be extended in various ways (Falconer and Mackay, 1996; Lynch and Walsh, 1998), with one of the most common being *P* = *A*+*PE*+*R*, where *PE* captures *dependent* effects, the so-called *permanent environmental effects*, while R captures the residual, *independent* variance that remains unexplained. Permanent environmental effects are stable differences among individuals above and beyond the permanent differences due to additive genetic effects. In repeated measurements of an individual, these effects create within-individual covariation. To prevent inflated estimates of additive genetic variance, these effects must therefore be modeled and estimated (Lynch and Walsh, 1998; Kruuk, 2004; Wilson et al., 2010).

This quantitative genetic decomposition of phenotypes is not possible at the individual level in non-clonal organisms, but under the crucial assumption of independence of genetic, permanent environmental, and residual effects, the phenotypic variance at the population level can be decomposed into the respective variance components as 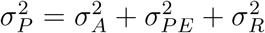. These variance components can then be used to understand and predict evolutionary change of phenotypic traits. For example, the additive genetic variance 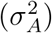 can be used to predict the response to selection using the breeder’s equation. It predicts the response to selection *R*_BE_ of a trait ***z*** (bold face notation denotes vectors) from the product of the heritability (*h*^2^) of the trait and the strength of selection (*S*) as

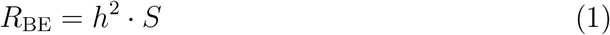

(Lush, 1937; Falconer and Mackay, 1996), where *h*^2^ is the proportion of additive genetic to total phenotypic variance

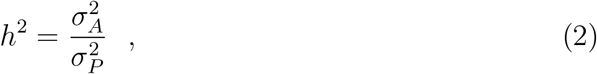

and *S* is the selection differential, defined as the mean phenotypic difference between selected individuals and the population mean or, equivalently, the phenotypic covariance *σ_p_*(***z, w***) between the trait (***z***) and relative fitness (***w***). Besides the breeder’s equation, evolution can be predicted using the secondary theorem of selection, according to which evolutionary change is equal to the additive genetic covariance of a trait with relative fitness, that is,

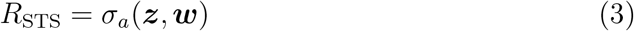

(Robertson, 1966; Price, 1970). Morrissey et al. (2010) and Morrissey et al. (2012) discuss the differences between the breeder’s equation and the secondary theorem of selection in detail. A major difference is that in contrast to *R*_BE_, *R*_STS_ only estimates the population evolutionary trajectory, but does not measure the role of selection in shaping this evolutionary change.

One measure of the role of selection is the selection gradient, which quantifies the strength of natural selection on a trait. For a normally distributed trait (***z***), it is given as the slope *β_z_* of the linear regression of relative fitness on a phenotypic trait (Lande and Arnold, 1983), that is,

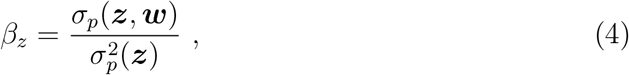

where 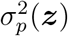 denotes the phenotypic variance of the trait, for which we only write 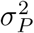 when there is no ambiguity about what trait the phenotypic variance refers to.

The reliable estimation of the parameters of interest (*h*^2^, *σ_p_*(***z, w***), *σ_a_*(***z, w***) and *β_z_*) and the successful prediction of evolution as *R_BE_* or *R_STS_*, require large amounts of data, often collected across multiple generations and with known relationships among individuals in the data set. For many phenotypic traits of interest, data collection is often not trivial, and multiple sources of error, such as phenotypic measurement error, pedigree errors (wrong relationships among individuals), or nonrandomly missing data may affect the parameter estimates. Several studies have discussed and addressed pedigree errors (*e.g*. Keller et al., 2001; Griffith et al., 2002; Senneke et al., 2004; Charmantier and Reale, 2005; Hadfield, 2008) and problems arising from missing data (*e.g*. Steinsland et al., 2014; Wolak and Reid, 2017). In contrast, although known for a long time (*e.g*. Price and Boag, 1987), the effects of phenotypic measurement error on estimates of (co-)variance components have received less attention (but see *e.g*. Hoffmann, 2000; Dohm, 2002; Macgregor et al., 2006; van der Sluis et al., 2010; Ge et al., 2017). In particular, general solutions to obtaining unbiased estimates of (co-)variance parameters in the presence of phenotypic measurement error are lacking.

In the simplest case, and the case considered here, phenotypic measurement error is assumed to be independent and additive, that is, instead of the actual phenotype ***z***, an error-prone version

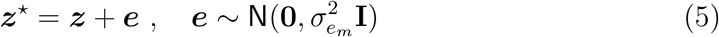

is measured, where ***e*** denotes an error term with independent correlation structure **I** and error variance 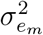 (see p.121 Lynch and Walsh, 1998). As a consequence, the *observed* phenotypic variance of the measured values is 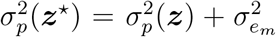, and thus larger than the *actual* phenotypic variance. The error variance a^p^_m_ thus must be disentangled from 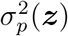 to obtain unbiased estimates of quantitative genetic parameters. However, most existing methods for continuous trait analyses that acknowledged measurement error have modeled it as part of the residual component, and thus implicitly as part of the total phenotypic value (*e.g*. Dohm, 2002; Macgregor et al., 2006; van der Sluis et al., 2010). This means that in the decomposition of a phenotype *P* = *A* + *PE* + *R*, measurement error is absorbed in *R*, thus 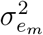 is absorbed by 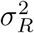. This practice effectively *downwardly* biases measures that are proportions of the phenotypic variance, in particular *h*^2^ and *β_z_*. To see why, let us denote the biased measures as 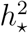 and 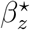. The biased version of heritability is then given as

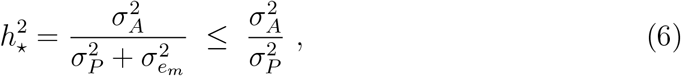

because under the assumption taken here that measurement error is independent of the actual trait value, measurement error is also independent of additive genetic differences and therefore leaves the estimate of the additive genetic variance 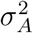 unaffected. This was already pointed out *e.g*. by Lynch and Walsh (p.121, 1998) or Ge et al. (2017). Equation (6) directly illustrates that 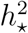 is attenuated by a factor 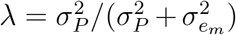, denoted as reliability ratio (*e.g*. Carroll et al., 2006). Using the same argument, one can show that 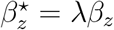, but also 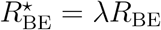, as will become clear later.

To obtain unbiased estimates of *h*^2^, *β_z_*, or any other quantity that depends on unbiased estimates of 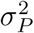, it is thus necessary to disentangle 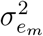 from the actual phenotypic variance 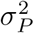, and particularly from its residual component 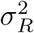. Importantly, however, purely mechanistic measurement imprecision is often not the only source of variation that may be considered irrelevant for the mechanisms of inheritance and selection in the system under study. Here, we therefore follow Ge et al. (2017) and use the term “transient effects” for the sum of measurement errors plus any biological short-term changes of the phenotype itself that are not considered relevant for the selection process, briefly denoted as “irrelevant fluctuations” of the actual trait.

As an example, if the trait is the mass of an adult animal, repeated measurements within the same day are expected to differ even in the absence of instrumental error, simply because animals eat, drink and defecate (for an example of the magnitude of these effects see Keller and Van Noordwijk, 1993). Such short-term fluctuations might not be of interest for the study of evolutionary dynamics, if the fluctuations do not contribute to the selection process in a given population. Under the assumption that these fluctuations are additive and independent among each other and of the actual trait value, they are mathematically indistinguishable from pure measurement error. In the remainder of the paper, we therefore do not introduce a separate notation to discriminate between (mechanistic) measurement error and biological short-term fluctuations, but treat them as a single component (***e***) with a total “error” variance 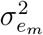. Consequently, we may sometimes refer to “measurement error” when in fact we mean transient effects as the sum of measurement error and transient fluctuations.

The aim of this article is to develop general methods to obtain unbiased estimates of heritability, selection, and response to selection in the presence of measurement error and irrelevant fluctuations of a trait, building on the work by Ge et al. (2017). We start by clarifying the meaning and information content of repeated phenotypic measurements on the same individual. The type of phenotypic trait we have in mind is a relatively plastic trait, such as milk production or an animal’s mass, which are expected to undergo changes across an individual’s lifespan that are relevant for selection. We show that repeated measures taken over different time intervals can help separate transient effects from more stable (permanent) environmental and genetic effects. We proceed to show that based on such a variance decomposition one can construct models that yield unbiased estimates of heritability, selection, and the response to selection. We illustrate these approaches with empirical quantitative trait analyses of body mass measurements taken in a population of snow voles in the Swiss alps (Bonnet et al., 2017).

## Material and methods

### Short-term and long-term repeated measurements

Table 1 gives an overview of how the different parameters considered here are (or are not) affected by the presence of measurement error. In order to retrieve unbiased estimates of all quantities given in Table 1, we must be able to appropriately model and estimate the measurement error variance 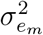, which can be achieved with repeated measurements. These repeated measurements must be taken in close temporal vicinity, that is, on a time scale where the focal trait is not actually undergoing any phenotypic changes that are relevant for selection. We introduce the notion of a *measurement session* for such *short-term* time intervals. In other words, a measurement session can be defined as a sufficiently short period of time during which the investigator is willing to assume that the residual component is constant. On the other hand, measurements are often repeated across much longer periods of time, such as months, seasons, or years, during which phenotypic change is not expected to be solely due to transient effects, and the resulting trait variation is often relevant for selection. Thus, *long-term* repeats, taken across different measurement sessions, help separating permanent environmental effects from residual components (*e.g*. Wilson et al., 2010).

**Table 1:** Overview of the effects of measurement error and transient fluctuations (ME) in a quantitative trait on important quantitative genetic parameters. The table indicates for each parameter whether it is biased or unbiased. For biased parameters the quantities are given that are estimated when ignoring transient effects in the quantitative genetic models. λ is the reliability ratio defined as 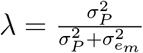. For notation see the main text.

| Parameter | Effect of ME | Biased parameter |
| --- | --- | --- |
| $\sigma_A^2$ | unbiased | - |
| $\sigma_{PE}^2$ | unbiased | - |
| $\sigma_R^2$ | biased | $\sigma_R^2 + \sigma_e^2$ |
| $h^2$ | biased | $\lambda h^2$ |
| $\beta_z$ | biased | $\lambda \beta_z$ |
| $\sigma_p(\mathbf{z}, \mathbf{w}) = S$ | unbiased | - |
| $\sigma_a(\mathbf{z}, \mathbf{w}) = R_{STS}$ | unbiased | - |
| $R_{BE}$ | biased | $\lambda R_{BE}$ |
| $I$ | unbiased | - |

The distinction between short-term and long-term repeats, and thus the definition of a measurement session, may not always be obvious or unique for a given trait. In the introduction we employed the example of an animal’s mass that transiently fluctuates within a day. Depending on the context, such fluctuations might not be of interest, and the “actual” phenotypic value would correspond to the average daily mass. A reasonable measurement session could then be a single day, and within-day repeats can thus be used to estimate 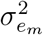. If however *any* fluctuations in body mass are of interest, irrespective of how persistent they are, much shorter measurement sessions, such as seconds or minutes, would be appropriate to ensure that only the purely mechanistic measurement error variance is represented by 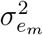.

### Repeated measurements in the animal model

In the following we show how measurement error can be incorporated in the key tool of quantitative genetics, the *animal model*, a special type of (generalized) linear mixed model, which is commonly used to decompose the phenotypic variance of a trait into genetic and non-genetic components (Henderson, 1976; Lynch and Walsh, 1998; Kruuk, 2004).

Let us assume that phenotypic measurements of a trait are blurred by measurement error following model (5), and that measurements have been taken both across and within multiple measurement sessions, as indicated in Figure 1a. Denoting by 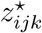 the *k*^th^ measurement of individual *i* in session *j*, it is possible to fit a model that decomposes the trait value as

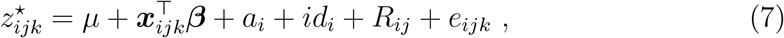

where *μ* is the population intercept, ***β*** is a vector of fixed effects and ***x***_*ijk*_ is the vector of covariates for measurement *k* in session *j* of animal *i*. The remaining components are the random effects, namely the breeding value *a_i_* with dependency structure 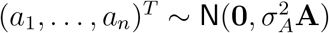, an independent, animal-specific permanent environmental effect 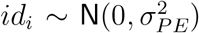, an independent Gaussian residual term 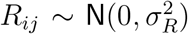, and an independent error term 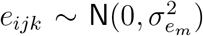 that absorbs any transient effects captured by the within-session repeats. The dependency structure of the breeding values *a_i_* is encoded by the additive genetic relatedness matrix **A** (Lynch and Walsh, 1998), which is traditionally derived from a pedigree, but can alternatively be calculated from genomic data (Meuwissen et al., 2001; Hill, 2014). The model can be further expanded to include more fixed or random effects, such as maternal, nest or time effects, but we omit such terms here without loss of generality. Importantly, model (7) does not require that all individuals have repeated measurements in each session in order to obtain an unbiased estimate of the variance components in the presence of measurement error. In fact, even if there are, on average, fewer than two repeated measurements per individual within sessions, it may be possible to separate the error variance from the residual variance, as long as the total number of within-session repeats over all individuals is reasonably large. We will in the following refer to model (7) as the “error-aware” model.

**Figure 1:**
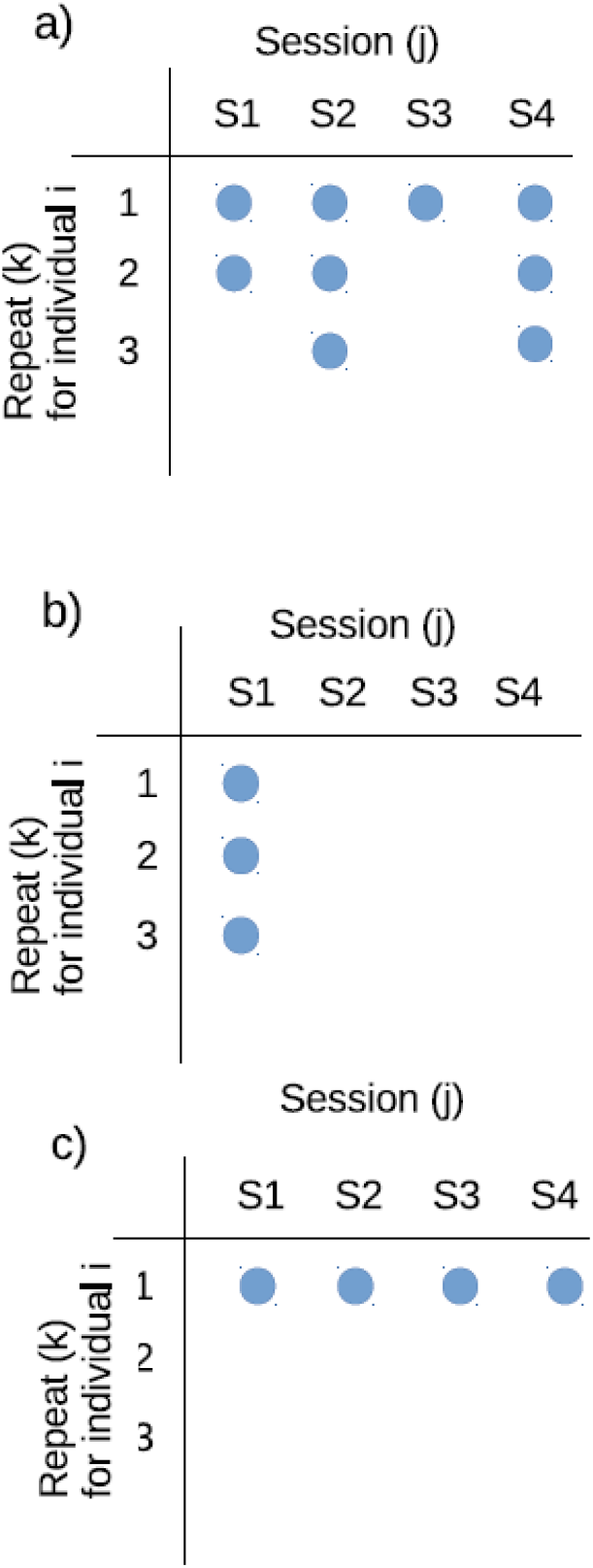
Schematic representation of three study designs, where one individual is measured a) multiple times across multiple measurement sessions, b) multiple times in one single measurement measurement session, or c) one single time across multiple measurement sessions. Only case a) allows to disentangle the measurement error variance 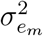 and the permanent environmental effects 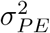 from 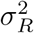, while case b) allows to separate only the measurement error variance and case c) only allows to disentangle permanent environmental effects.

If, however, a trait has not been measured across different time scales (*i.e*. either only within or only across measurement sessions), not all variance components are estimable. In the first case, when repeats are only taken within a single measurement session for each individual, as depicted in Figure 1b, an error term can be included in the model, but a permanent environmental effect cannot. The model must then be reduced to

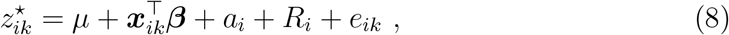

thus it is possible to estimate the error variance 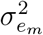 and to obtain unbiased estimates of 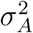 and *h*^2^, while the residual variance 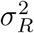 then also contains the permanent environmental variance. In the second case, when repeated measurements are only available from across different measurement sessions, as illustrated in Figure 1c, the error variance cannot be estimated. Instead, an animal-specific permanent environmental effect can be added to the model, which is then given as

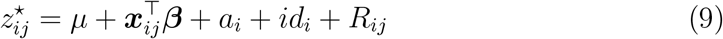

for the measurement in session *j* for individual *i*. Interestingly, this last model mirrors the types of repeats that motivated quantitative geneticists to isolate 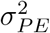, which may otherwise be confounded not only with 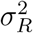, but also with 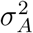. This occurs because the repeated measurements across sessions induce an increased within-animal correlation (*i.e*. a similarity) that may be absorbed by 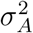 if not modeled appropriately (Kruuk and Hadfield, 2007; Wilson et al., 2010).

### Measurement error and selection

Selection occurs when a trait is correlated with fitness, such that variations in the trait values lead to predictable variations among the same individuals in fitness. The leading approach for measuring the strength of directional selection is the one developed by Lande and Arnold (1983), who proposed to estimate the selection gradient *β_z_* as the slope of the regression of relative fitness ***w*** on the phenotypic trait ***z***

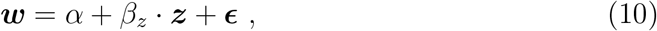

with intercept a and residual error vector ***ϵ***. This model can be further extended to account for covariates, such as sex or age. If the phenotype ***z*** is measured with error (which may again encompass any irrelevant fluctuations), such that the observed value is ***z***^⋆^ = ***z*** + ***e*** with error variance 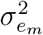 as in (5), the regression of ***w*** against ***z***^⋆^ leads to an attenuated version of *β_z_* (Mitchell-Olds and Shaw, 1987; Fuller, 1987; Carroll et al., 2006). Using that 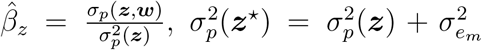, and the assumption that the error in ***z***^⋆^ is independent of ***w***, simple calculations show that the error-prone estimate of selection is

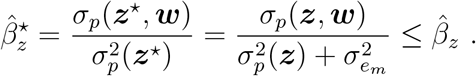

Hence, the quantity that is estimated is 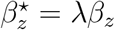 with 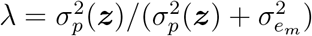, thus *β_z_* suffers from exactly the same bias as the estimate of heritability (see again Table 1). To obtain an unbiased estimate of selection it may thus often be necessary to account for the error by a suitable error model. Such error-aware model must rely on the same type of short-term repeated measurements as those used in (7) or (8), but with the additional complication that ***z*** is now a covariate in a regression model, and no longer the response. In order to estimate an unbiased version of *β_z_* we therefore rely on the interpretation as an error-in-variables problem for classical measurement error (Fuller, 1987; Carroll et al., 2006). To this end, we propose to formulate a *Bayesian hierarchical model*, because this formulation, together with the possibility to include prior knowledge, provides a flexible way to model measurement error (Stephens and Dellaportas, 1992; Richardson and Gilks, 1993). To obtain an error-aware model that accounts for error in selection gradients, we need a three-level hierarchical model: The first level is the regression model for selection, and the second level is given by the error model of the observed covariate ***z***^⋆^ given its true value ***z***. Third, a so-called *exposure model* for the unobserved (true) trait value is required to inform the model about the distribution of ***z***, and it seems natural to employ the animal model (9) for this purpose. Again using the notation for an individual *i* measured in different sessions *j* and with repeats *k* within sessions, the formulation of the three-level hierarchical model is given as

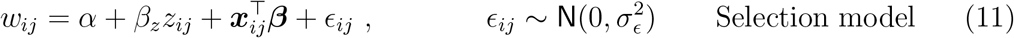

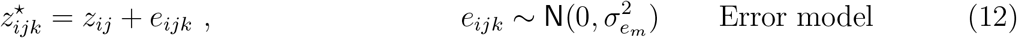

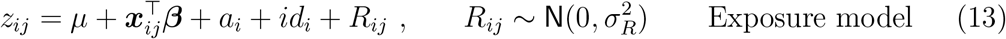

where *w_ij_* is the measurement of relative fitness for individual *i*, usually taken only once per individual and having the same value for all measurement sessions *j*, ***β*** is a vector of fixed effects, ***x***_*ij*_ is the vector of covariates for animal *i* in measurement session *j, β_z_* is the selection gradient, and *α* and *ϵ_ij_* are respectively the intercept and the independent residual term from the linear regression model. The classical independent measurement error term is given by *e_ijk_*. This formulation as a hierarchical model gives an unbiased estimate of the selection gradient *β_z_*, because the lower levels of the model properly account for the error in z by explicitly modelling it. It might be helpful to see that the second and third levels are just a hierarchical representation of model (7). Model (11)-(13) can be fitted in a Bayesian setup, see for instance Muff et al. (2015) for a description of the implementation in INLA (Rue et al., 2009) via its `R` interface `R-INLA`.

Note that model (11) is formulated here for directional selection. Although the explicit discussion of alternative selection mechanisms, such as stabilizing or disruptive selection, is beyond the scope of the present paper, we note that error modelling for these cases is straightforward: The only change is that the linear selection model (11) is replaced by the appropriate alternative, for example by including quadratic or any other kind of non-linear terms (*e.g*. Fisher, 1930; Lande and Arnold, 1983). Moreover, (11) can be replaced by any other regression model, for example by one that accounts for non-normality of fitness (see *e.g*. Morrissey and Sakrejda, 2013; Morrissey and Goudie, 2016). Similarly, it is conceptually straightforward to replace the Gaussian error and exposure models, if there is reason to believe that the normal assumptions for the error term *e_ijk_* or the residual term *R_ij_* are unrealistic, for example if ***z*** is a count or a binary variable. In fact, equation (10) to estimate selection does not actually assume a specific distribution for ***z***, however the interpretation of *β_z_* as a directional selection gradient to predict evolutionary change may be lost for non-Gaussian traits (Lande and Arnold, 1983). Finally and importantly, although multivariate selection is not covered in the present paper, it is possible to extend the hierarchical model (11)-(13) to the multivariate case.

### Measurement error and the response to selection

#### The breeder’s equation

Evolutionary response to a selection process on a phenotypic trait can be predicted either by the breeder’s equation (1) or by the Robertson-Price identity (3), and these two approaches are equivalent only when the respective trait value (in the univariate model) is the sole causal factor affecting fitness (Morrissey et al., 2010, 2012). Even if the breeder’s equation is formulated for multiple traits, the implicit assumption still is that *all* correlated traits causally related to fitness are included in the model. Given that fitness is a complex trait that usually depends on many unmeasured variables (Møller and Jennions, 2002; Peek et al., 2003), it is not surprising that the breeder’s equation is often not successful in predicting evolutionary change in natural systems (Hadfield, 2008; Morrissey et al., 2010), in contrast to (artificial) animal breeding situations, where, thanks to the control over the process, all the traits affecting fitness are known and included in the models (Lush, 1937; Falconer and Mackay, 1996; Roff, 2007).

To understand how transient effects affect the estimate of *R_BE_* = *h*^2^ · *S*, we must understand how the components *h*^2^ and *S* are affected. We have seen that 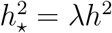. On the other hand, the selection differential *S*_⋆_ = *σ_p_*(***z^⋆^, w***) is an unbiased estimate of *σ_p_*(***z, w***), because under the assumption of independence of the error vector ***e*** and fitness ***w***,

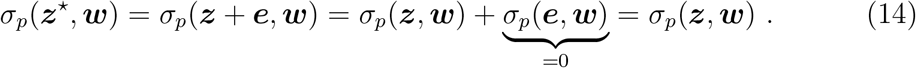

Consequently, the bias in 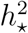 directly propagates to the estimated response to selection, that is, 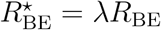 (Table 1).

#### The Robertson-Price identity

Response to selection can also be predicted using the secondary theorem of selection. Specifically, the additive genetic covariance of the relative fitness ***w*** and the phenotypic trait ***z***, *σ_a_*(***w, z***) can be estimated from a bivariate animal model. If interest centers around the evolutionary response of a single trait, the model for the response vector including the (error-prone) trait values ***z***^⋆^ and relative fitness values ***w*** is bivariate with

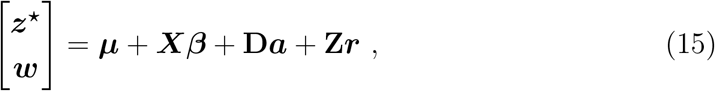

where ***μ*** is the intercept vector, ***β*** the vector of fixed effects, ***X*** the corresponding design matrix, **D** is the design matrix for the breeding values ***a***, and **Z** is a design matrix for additional random terms ***r***. These may include environmental and/or error terms, depending on the structure of the data, that may correspond to the univariate cases of equations (7) – (9) or again to other random terms such as maternal or nest effects. The actual component of interest is the vector of breeding values, which is assumed multivariate normally distributed with

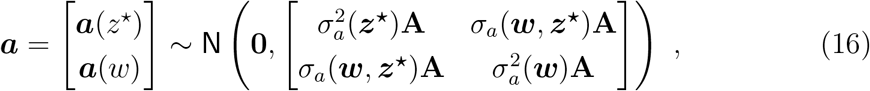

where ***a***(*z*^⋆^) and ***a***(*w*) are the respective subvectors for the trait and fitness, and **A** is the relationship matrix derived from the pedigree. An estimate of the additive genetic covariance *σ_a_*(***w, z***^⋆^) is extracted from this covariance matrix. An interesting feature of the additive genetic covariance, and consequently estimates of the response to selection using the STS, is that it is unbiased by independent error in the phenotype. This can be seen by reiterating the exact same argument as in equation (14), but replacing the phenotypic with the genetic covariance.

We confirmed all these theoretical expectations with a simulation study, where we analysed the effects of measurement error on the estimates of interest by adding error terms with different variances to the phenotypic traits. Details and results of the simulations are given in Appendix 2, while the code for their implementation is reported in Appendix 3.

### Example: Body mass of snow voles

The empirical data we use here stem from a snow vole population that has been monitored between 2006 and 2014 in the Swiss Alps (Bonnet et al., 2017). The genetic pedigree is available for 937 voles, together with measurements on morphological and life history traits. Thanks to the isolated location, it was possible to monitor the whole population and to obtain high recapture probabilities (0.924 ± 0.012 for adults and 0.814 ± 0.030 for juveniles). Details of the study are given in Bonnet et al. (2017).

Our analyses focused on the estimation of quantitative genetic parameters for the animals’ body mass (in grams). The dataset contained 3266 mass observations from 917 different voles across 9 years. Such measurements are expected to suffer from classical measurement error, as they were taken with a spring scale, which is prone to measurement error under field conditions. In addition, the actual mass of an animal may contain irrelevant within-day fluctuations (eating, defecating, digestive processes), but also unknown pregnancy conditions in females, which cannot reliably be determined in the field. Repeated measurements were available, both recorded within and across different seasons. In each season two to five “trapping sessions” were conducted, which each lasted four consecutive nights. Although this definition of measurement session was based purely on operational aspects driven by the data collection process, we used this time interval to estimate 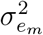. It is arguably possible that four days might be undesirably long, and that variability in such an interval includes more than purely transient effects, but the data did not allow for a finer time-resolution. However, to illustrate the importance of the measurement session length, we also repeated all analyses with measurement sessions defined as a calendar month, which is expected to identify a larger (and probably too high) proportion of variance as 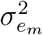. The number of 4-day measurement sessions per individual was on average 3.02 (min = 1, max = 24) with 1.15 (min = 1, max = 3) number of shortterm repeats on average, while there were 2.37 (min = 1, max = 13) one-month measurement sessions on average, with 1.41 (min = 1, max = 6) short-term repeats per measurement session.

#### Heritability

Bonnet et al. (2017) estimated heritability using an animal model with sex, age, Julian date (JD), squared Julian date and the two-way and three-way interactions among sex, age and Julian date as fixed effects. The inbreeding coefficient was included to avoid bias in the estimation of additive genetic variances (de Boer and Hoeschele, 1993). The breeding value (*a_i_*), the maternal identity (*m_i_*) and the permanent environmental effect explained by the individual identity (*id_i_*) were included as individual-specific random effects.

If no distinction is made between short-term (within measurement session) and long-term (across measurement sessions) repeated measurements, the model that we denote as the *naive* model is given as

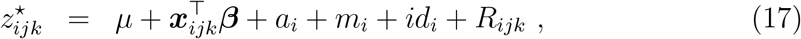

where 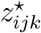 is the mass of animal *i* in measurement session *j* for repeat *k*. This model is prone to underestimate heritability, because it does not separate the variance 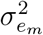 from the residual variability, and 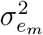 is thus treated as part of the total phenotypic trait variability. To isolate the measurement error variance, the model expansion

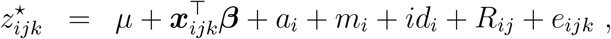

with 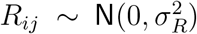 and 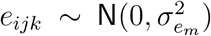 leads to what we denote here as the *error-aware* model. Under the assumption that the length of a measurement session was defined in an appropriate way, and that the error obeys model (5), this model yields an unbiased estimate of *h*^2^, calculated as 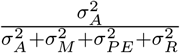 (in agreement with Bonnet et al., 2017), where 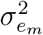 is explicitly estimated and thus not included in the denominator. Both models were implemented in `MCMCglmm` and are reported in Appendix 4. Inverse gamma priors `IG`(0.01,0.01), parameterized with shape and rate parameters, were used for all variances in all models, while `N`(0,10^12^) (*i.e*. default `MCMCglmm`) priors were given to the fixed effect parameters. Analyses were repeated with varying priors on 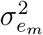 for a sensitivity check, but results were very robust (results not shown).

#### Selection

Selection gradients were estimated from the regression of relative fitness (***w***) on body mass (***z***^⋆^). Relative fitness was defined as the relative lifetime reproductive success (rLRS), calculated as the number of offspring over the lifetime of an individual, divided by the population mean LRS. The naive estimate of the selection gradient was obtained from a linear mixed model (*i.e*. treating rLRS as continuous trait), where body mass, sex and age were included as fixed effects, plus a cohort-specific random effect. The error-aware version of the selection gradient *β_z_* was estimated using a three-layer hierarchical error model as in (11)-(13) that also included an additional random effect for cohort in the regression model. Sex and age were also included as fixed effects in the exposure model, plus breeding values, permanent environmental and a residual term as random effects. The hierarchical model used to estimate the error-aware *β_z_* was implemented in `INLA` and is described in Appendix 1, with `R` code given in Appendix 5. Again, `IG`(0.01, 0.01) priors were assigned to all variance components, while independent `N`(0,10^2^) priors were used for all slope parameters. Since rLRS is not actually a Gaussian trait, *p*-values and CIs of the estimate for *β_z_* from the linear regression model are, however, incorrect. Although recent considerations indicate that selection gradients could directly be extracted from an overdispersed Poisson model (Morrissey and Goudie, 2016), we followed the original analysis of Bonnet et al. (2017) and extracted *p*-values from an overdispersed Poisson regression model with absolute LRS as a count outcome, both for the (naive) model without error modelling *and* for the hierarchical error model, where the linear model (13) was replaced by an overdispersed Poisson regression model (see Appendices 1 and 5 for the model description and code for both models).

#### Response to selection

Response to selection on body mass was estimated with rLRS using the breeder’s equation (1) and the secondary theorem of selection (3), both for the naive and the error-aware versions of the model. The naive and error-aware versions of *R*_BE_ were estimated by substituting either the naive 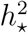 or the error-aware estimates of *h*^2^ into the breeder’s equation, where the selection differential was calculated as the phenotypic covariance between mass and rLRS. On the other hand, *R*_STS_ was estimated from the bivariate animal model, implemented in `MCMCglmm` using the same fixed and random effects as those in equation (17). Again `IG`(0.01, 0.01) priors were used for the variance components. No residual component was included for the fitness trait, as suggested by Morrissey et al. (2012), and its error variance was fixed at 0, because no error modelling is required. Appendix 6 contains the respective `R` code.

## Results

### Heritability

As expected from theory (Table 1), transient effects in the measurements of body mass biased some, but not all, quantitative genetic estimates in our snow vole example (Table 2). The estimates and confidence intervals of the additive genetic variance 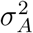, as well as the permanent environmental variance 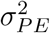 and the maternal variance (denoted as 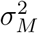) were only slightly corrected in the error-aware models. Residual variances, however, were much lower when measurement error was accounted for in the models. The measurement error model separated residual and transient (error) variance so that 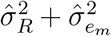 corresponded approximately to 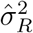 from the naive model. The overestimation of the residual variance resulted in estimates of heritability that were underestimated by nearly 40% when measurement error was ignored (*ĥ*^2^ = 0.14 in the naive model and *ĥ*^2^ = 0.23 in the error-aware model).

**Table 2:** Estimates of quantitative genetic parameters of body mass in snow voles using naive and error-aware models. The posterior modes of variance components and heritability are given, together with their 95% credible intervals (in brackets).

| model | $\hat{h}^2$ | $\hat{\sigma}_A^2$ | $\hat{\sigma}_{PE}^2$ | $\hat{\sigma}_M^2$ | $\hat{\sigma}_R^2$ | $\hat{\sigma}_{em}^2$ |
| --- | --- | --- | --- | --- | --- | --- |
| naive | 0.14<br>[0.07, 0.25] | 3.40<br>[1.41, 6.15] | 6.09<br>[4.33, 8.51] | 1.16<br>[0.56, 2.84] | 12.40<br>[11.78, 13.21] | - |
| error-aware<br>(4-day measurement session) | 0.23<br>[0.09, 0.33] | 3.97<br>[1.46, 6.06] | 5.62<br>[3.68, 7.68] | 1.48<br>[0.57, 2.73] | 6.58<br>[5.76, 7.82] | 6.07<br>[5.54, 7.05] |
| error-aware<br>(one-month measurement session) | 0.24<br>[0.10, 0.37] | 3.82<br>[1.17, 5.84] | 4.78<br>[3.16, 7.21] | 1.58<br>[0.61, 2.86] | 5.77<br>[4.78, 6.71] | 7.91<br>[7.15, 8.38] |

As expected, the estimated measurement error variance is larger when a measurement session is defined as a full month 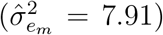 than as a 4-day interval (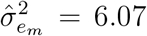, Table 2), because the trait then has more time and opportunity to change. As a consequence, heritability is even slightly higher (*ĥ*^2^ = 0.24) when the longer measurement session definition is used. This example is instructive because it underlines the importance of defining the time scale at which short-term repeats are expected to capture only transient, and not biologically relevant variability of the phenotypic trait. In the case of the mass of a snow vole, most biologists would probably agree that changes in body mass over a one-month measurement session may well be biologically meaningful (*i.e*. body fat accumulation, pregnancy in females, etc.), while it is less clear how much of the fluctuations within a 4-day measurement session are transient, and what part of it would be relevant for selection. Within-day repeats might be the most appropriate for the case of mass, since within-day variance is likely mostly transient, but because the data were not collected with the intention to quantify such effects, within-day repeats were not available in sufficient numbers in our example data set.

### Selection

As expected, estimates of selection gradients 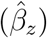 obtained with the measurement error models provided nearly 40% higher estimates of selection than the naive model (Table 3). The two measurement session lengths yielded similar results. With and without measurement error modelling, the *p*-values of the zero-inflated Poisson models confirmed the presence of selection on body mass in snow voles (*p* < 0.001 in all models).

**Table 3:** Estimates of selection gradients 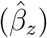 for body mass in snow voles, derived from naive (ML estimate) and error-aware models (posterior means). For both types of models, Bayesian *p*-values were derived from zero-inflated Poisson regressions.

| model | $\hat{\beta}_z$ | $p$-value |
| --- | --- | --- |
| naïve | 0.065 | < 0.001 |
| error-aware<br>(4-day measurement session) | 0.104 | < 0.001 |
| error-aware<br>(one-month measurement session) | 0.104 | < 0.001 |

### Response to selection

In line with theory, estimates of the response to selection using the breeder’s equation were nearly 40% higher when transient effects were incorporated in the quantitative genetic models using 4-day measurement sessions (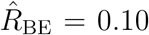 in the naive model and 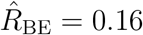 in the error-aware model; Table 4). As in the case of heritability, the one-month measurement session definition resulted in even slightly higher estimates of the response to selection 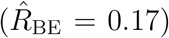. In contrast, response to selection measured by the secondary theorem of selection 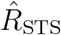 did not show evidence of bias, and the error-aware model with a 4-day measurement session definition estimated the same value 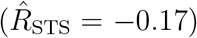 as the naive model (Table 4). With a one-month measurement session, we obtained a slightly attenuated value 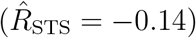, although the difference was small in comparison to the credible intervals (Table 4).

**Table 4:** Response to selection for body mass in snow voles (posterior modes and 95% credible intervals) estimated with the breeder’s equation 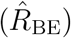 and with the secondary theorem of selection 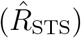. Results are shown for the naive and the error-aware models.

| model | $\hat{R}_{\text{STS}}$ | 95% CI | $\hat{R}_{\text{BE}}$ | 95% CI |
| --- | --- | --- | --- | --- |
| naive | -0.17 | $[-0.54, 0.18]$ | 0.10 | $[0.05, 0.17]$ |
| error-aware<br>(4-day measurement session) | -0.17 | $[-0.51, 0.19]$ | 0.16 | $[0.06, 0.23]$ |
| error-aware<br>(one-month measurement session) | -0.14 | $[-0.53, 0.17]$ | 0.17 | $[0.07, 0.26]$ |

This example illustrates that the breeder’s equation is generally prone to underestimation of the selection response in real study systems when measurement error in the phenotype is present (Table 1). The results also confirm that estimates for response to selection may differ dramatically between the breeder’s equation and the secondary theorem of selection approach. As already noticed by Bonnet et al. (2017), the predicted evolutionary response derived from the breeder’s equation points in the opposite direction in the snow vole data than the estimate derived from the secondary theorem of selection (*e.g*. naive estimates 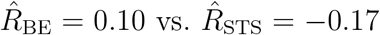, with non-overlapping credible intervals; Table 4).

## Discussion

This study addresses the problem of measurement error and transient fluctuations in continuous phenotypic traits in quantitative genetic analyses. We show that measurement error and transient fluctuations can lead to substantial bias in estimates of several important quantitative genetic parameters, including heritability, selection gradients and the response to selection (Table 1). We introduce modelling strategies to obtain unbiased estimates in these parameters in the presence of measurement error and transient fluctuations. These strategies rely on the distinction between variability from stable effects that are part of the biologically relevant phenotypic variability, and transient effects, which are the sum of mechanistic measurement error and biological fluctuations that are considered irrelevant for the selection process. We argue that ignoring the distinction between stable and transient effects may not only lead to an underestimation of the heritability due to inflated estimates of the residual variance, aR, but also to bias in the estimates of selection gradients and the response to selection. Measurements of the same individual repeated at appropriate time scales allow the variance from such transient effects to be partitioned, and thus prevent such bias.

How can repeated measurements be used to prevent an *under*estimation of heritability, selection, and response to selection, while permanent environment effects are required in quantitative genetic models of repeated measures to avoid an upward bias of 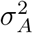 and, hence, an over estimation of *h*^2^ (Wilson et al., 2010)? The fact that repeated measurements are used to prevent opposite biases in heritability estimates makes it apparent that the information content in what is termed “repeated measurements” in both cases is very different. The crucial aspect is that it matters at which temporal distance the repeats were taken, and that the relevance of this distance depends on the kind of trait under study. Repeats taken on the same individual at different life stages (“long-term” repeats, *e.g*. across what we call measurement sessions here) can be used to separate the animal-specific permanent environmental effect from both genetic and residual variances. On the other hand, repeats taken in temporal vicinity (“short-term” repeats, *e.g*. within a measurement session) help disentangle any transient from the residual effects. Only by modelling *both* types of repeats, that is, across different relevant time scales, is it practically feasible to separate all variance components. To do so, the quantitative genetic model for the trait value, typically the animal model, needs extension to three levels of measurement hierarchy (equation (7)): the individual (*i*), the measurement session (*j* within *i*) and the repeat (*k* within *j* within *i*). As highlighted with the snow vole example, it may not always be trivial to determine, in a particular system, an appropriate distinction between short-term and long-term repeats, and consequently how to define a measurement session. This decision must be driven by the definition of short-term variation as a variation that is not “seen” by the selection process (see *e.g*. Price and Boag, 1987, p. 279 for a similar analogy), in contrast to persistent effects that are potentially under selection. This distinction ultimately depends on the trait, on the system under study and on the research question that is asked, because some traits may fluctuate on extremely short time scales (minutes or days), while others remain constant across an entire adult’s life.

The application to the snow vole data, where we varied the measurement session length from four days to one month, illustrated that longer measurement sessions automatically capture more variability, that is, the estimated error variance 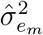 increased. Consequently, unreasonably long measurement sessions may lead to overcorrected estimates of the parameters of interest. On the other hand, considering measurement sessions that are too short may lead to an insufficient number of within-session repeats, or they may fail to identify transient variability that is biologically irrelevant. This makes clear that a careful definition of measurement session length is important already at the design stage of a study.

If one is uncertain whether repeated measurements capture effects relevant to selection or not, would averaging over repeats result in better estimates of quantitative genetic measures? Averaging methods have been proposed specifically to reduce bias that emerges due to measurement error and transient effects (Carbonaro et al., 2009; Zheng et al., 2016). While averaging will alleviate bias by reducing the error variance in the mean, it will not eliminate it completely. This can be seen from the fact that averaging over *K* within-session repeats for all animals and measurement sessions, the variance 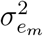 is reduced to 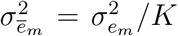, assuming independence of the error term. Unless *K* is large, 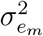 will not approach zero. Moreover, this practice only works if all animals have the same number of repeats within all measurement sessions, but it will not work in the unbalanced sampling design so common in studies of natural populations.

Our method approaches the problem of measurement error and transient fluctuations by assuming a dichotomous distinction between short-term and long-term repeats. An alternative perspective of within-animal repeated measurements could take a continuous view, recalling that repeated measurements are usually correlated, even when taken across long time spans, and that the correlation increases the closer in time the measurements were taken. A more sophisticated model could thus take into account that the residual component in the model changes continuously, and introduce a time-dependent correlation structure instead of simply distinguishing between short-term and long-term repeats. Such a model might be beneficial if repeats were not taken in clearly defined measurement sessions, although such a temporal correlation term introduces another level of model complexity, and thus entails other challenges.

It may sometimes not be possible to take multiple measurements on the same individual, or to repeat a measurement within a session. However, it may still be feasible to include an appropriate random effect in the absence of short-term repeats, provided that knowledge about the error variance is available, *e.g*. from previous studies that used the same measurement devices, from a subset of the data, or from other “expert” knowledge. The Bayesian framework is ideal in this regard, because it is straightforward to include random effects with a very strong (or even fixed) prior on the respective variance component. Such Bayesian models provide error-aware estimates that are equivalent to those illustrated in Table 1, but with the additional advantage that posterior distributions naturally reflect all uncertainty that is present in the parameters, including the uncertainty that is incorporated in the prior distribution of the error variance.

Measurement error and transient fluctuations bias some, but not all quantitative genetic inferences. When 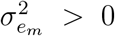, the naive estimates of *h*^2^, *β_z_* and *R*_BE_ are attenuated by the same factor λ < 1, but other components, such as the selection differential *S* or *R*_STS_, are not affected (Table 1). The robustness of the secondary theorem of selection to measurement error can certainly be seen as an advantage over the breeder’s equation. Nevertheless, the Robertson-Price identity does not model selection explicitly, and thus says little about the selective processes. The Robertson-Price equation can be used to check the consistency of predictions made from the breeder’s equation, but the breeder’s equation remains necessary to test hypothesis about the causal nature of selection (Morrissey et al., 2012; Bonnet et al., 2017). Another quantity that is unaffected by independent transient effects, which we however did not further elaborate on here, is *evolvability*, defined as the squared coefficient of variation 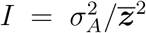, where 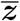 denotes the mean phenotypic value (Houle, 1992). Evolvability is often used as an alternative to heritability, and is interpreted as the *opportunity for selection* (Crow, 1958). Not only 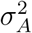, but also 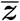 can be consistently estimated using ***z***^⋆^, namely because the expected values `E`[***z***^⋆^] = `E`[***z***] due to the independence and zero mean of the error term. For completeness, we added evolvability to Table 1.

A critical assumption of our models was that the error components are independent of the phenotypic trait under study, but also independent of fitness or any covariates in the animal model or the selection model. While the small changes in *R*_STS_ that we observed in the snow vole application with one-month measurement sessions could be due to pure estimation stochasticity, an alternative interpretation is that the measurement error in the data are not independent of the animal’s fitness. At least two processes could lead to a correlation between the measurement error in mass and fitness in snow voles. First, pregnant females will experience temporally increased body mass, and we expect the positive deviation from the true body mass to be correlated with fitness, because a pregnant animal is likely to have a higher expected number of offspring over its entire lifespan. And second, some of the snow voles were not fully grown when measured, and juveniles are more likely to survive if they keep growing, so that deviations from mean mass over the measurement session period would be non-randomly associated with life-time fitness.

So far, we have focused on traits that can change relatively quickly throughout the life of an individual, such as body mass, or physiological and behavioral traits. Traits that remain constant after a certain age facilitate the isolation of measurement error, because the residual variance term is then indistinguishable from the error term, given that a permanent environmental (*i.e*. individual-specific) effect is included in the model. In such a situation it is sufficient to estimate 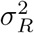, which then automatically corresponds to the measurement error variance, while 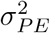 captures all the environmental variability. However, not many traits will fit that description. The majority of traits, even seemingly stable traits such as skeletal traits, are in fact variable over time (Price and Grant, 1984; Smith et al., 1986).

We have shown that dealing appropriately with measurement error and transient fluctuations of phenotypic traits in quantitative genetic analyses requires the inclusion of additional variance components. Quantitative genetic analyses often differ in the variance components that are included to account for important dependencies in the data (Meffert et al., 2002; Palucci et al., 2007; Kruuk and Hadfield, 2007; Hadfield et al., 2013). Besides the importance of separating the right variance components, it has been widely discussed which of the components are to be included in the denominator of heritability estimates, although the focus has been mainly on the proper handling of variances that are captured by the fixed effects (Wilson, 2008; de Villemereuil et al., 2018). We hope that our treatment of measurement error in quantitative genetic analyses sparks new discussions of what should be included in the denominator when heritability is calculated.

The methods presented in this paper have been developed and implemented for continuous phenotypic traits. Binary, categorical or count traits may also suffer from measurement error, which is then denoted as misclassification error (Copas, 1988; Magder and Hughes, 1997; Küchenhoff et al., 2006), or as miscounting error (*e.g*. Muff et al., 2018). Models for non-Gaussian traits are usually formulated in a generalized linear model framework (Nakagawa and Schielzeth, 2010; de Villemereuil et al., 2016) and require the use of a link function (*e.g*. the logistic or log link). In these cases, it will often not be possible to obtain unbiased estimates of quantitative genetic parameters by adding an error term to the linear predictor as we have done here for continuous traits. Obtaining unbiased estimates of quantitative genetic parameters in the presence of misclassification and miscounting error will require extended modelling strategies, such as hierarchical models with an explicit level for the error process.

We hope that the concepts and methods provided here serve as a useful starting point when estimating quantitative genetics parameters in the presence of measurement error or transient, irrelevant fluctuations in phenotypic traits. The proposed approaches are relatively straightforward to implement, but further generalizations are possible and will hopefully follow in the future.

## Supporting information

**Appendix 1:** Supplementary text and figures (pdf)

**Appendix 2:** Supplementary text and figures for simulation study (pdf)

**Appendix 3:** R script for the simulation and analysis of pedigree data

**Appendix 4:** R script for heritability in snow voles

**Appendix 5:** R script for selection in snow voles

**Appendix 6:** R script for response to selection in snow voles.

